# A custom library construction method for super-resolution ribosome profiling in Arabidopsis

**DOI:** 10.1101/2022.07.14.499987

**Authors:** Hsin-Yen Larry Wu, Polly Yingshan Hsu

## Abstract

Ribosome profiling (aka Ribo-seq) is the deep sequencing of ribosome footprints (RFs). It maps and quantifies ribosome occupancy on mRNA, which enables the identification of coding regions and the accurate quantification of translation efficiency. We previously optimized the Ribo-seq method in Arabidopsis and tomato (Hsu et al., 2016; Wu et al., 2019; Wu and Hsu, 2022) to obtain precise RFs with strong 3-nucleotide periodicity, a feature displayed by actively translating ribosomes and a benchmark of high-quality Ribo-seq (Brar and Weissman, 2015). This strong periodicity allowed us to confidently define numerous unannotated translation events across plants (Hsu et al., 2016; Wu et al., 2019; Wu and Hsu, 2022). Recently, several key commercial reagents used in our methods were discontinued; thus, there is an urgent need to develop a new protocol. Here, we report an updated protocol that adapts two custom library construction methods (McGlincy and Ingolia, 2017; Li et al., 2021) for plants. We applied this new protocol to Arabidopsis seedlings and obtained high-quality data. We describe our step-by-step method and discuss crucial considerations for Ribo-seq experiments. We also provide a bioinformatic pipeline to perform essential quality control analyses on Ribo-seq data. Our approach should be readily applicable to other plant species with minimal modifications.

## Results and Discussion

### Considerations for Ribo-seq experiments

Ribo-seq experiments can be divided into two phases: ribosome footprinting and sequencing library construction. Ribosome footprinting involves 1) lysate preparation, 2) ribonuclease digestion, 3) monosome isolation and RNA purification, and 4) RF size selection. In addition, rRNA depletion is commonly deployed to eliminate abundant rRNA contaminants during library construction (**Figure 1A**).

**Figure 1.**
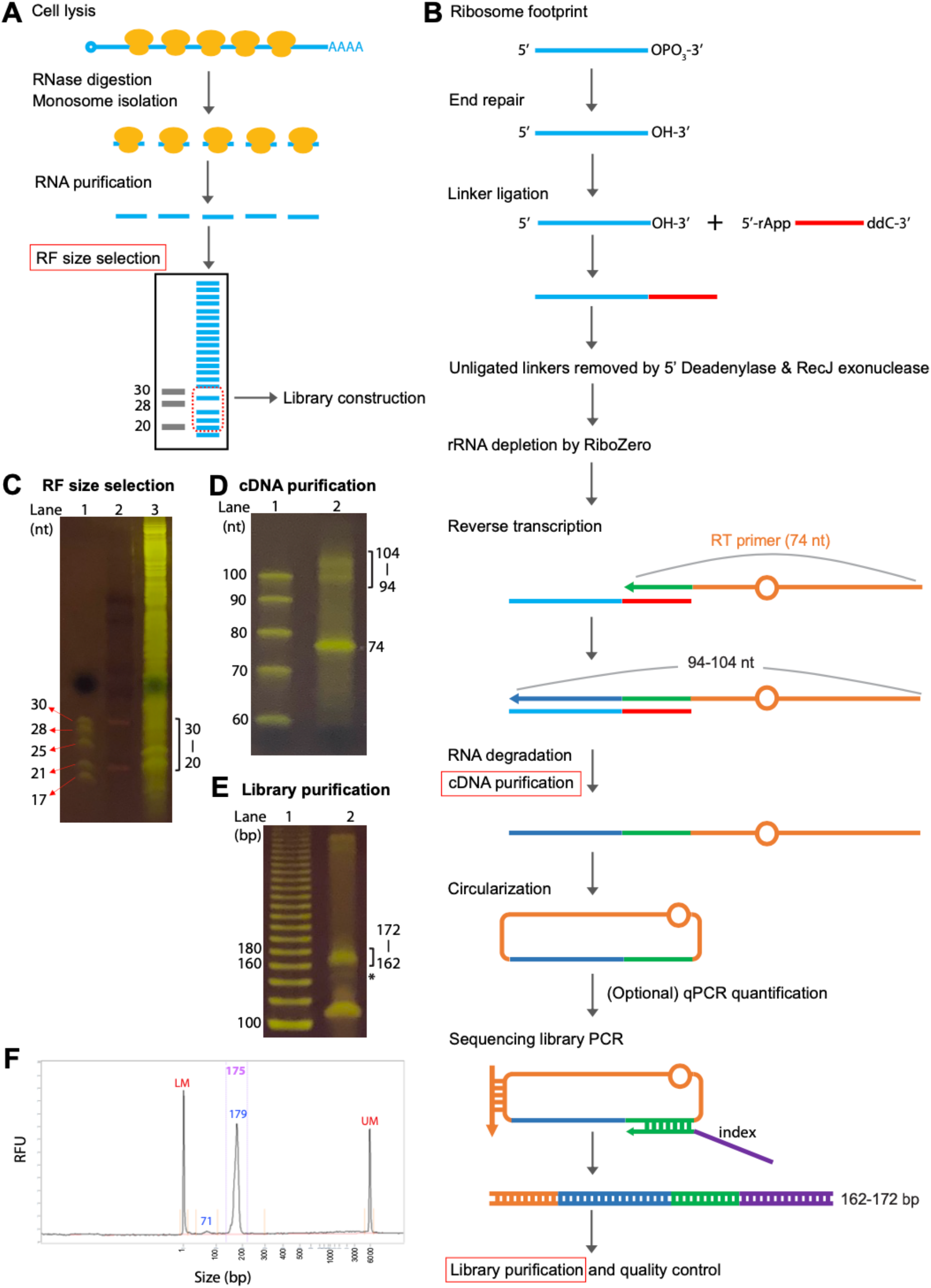
Ribo-seq workflow and representative gel images. **(A)** Ribo-seq overview: A Ribo-seq experiment consists of ribosome footprinting and sequencing library construction. **(B)** Workflow for custom Ribo-seq library construction. After size selection, ribosome footprints undergo end repair, ligation to a linker, removal of excess linkers, rRNA depletion, reverse transcription, cDNA purification, cDNA circularization, library PCR, and library purification. Among these steps, three (size selection of ribosome footprints, cDNA purification, and library purification) involve gel purifications and are highlighted by red boxes. For 20-30-nt ribosome footprints, the expected cDNA length would be 94-104 nt, and the expected library size would be 162-172 bp. **(C)** Size selection of ribosome footprints using a 15% TBE-urea gel. Lane 1: 30- and 28-nt marker (from the discontinued illumina Ribo-seq kit) and 25-, 21-, 17-nt marker (NEB microRNA marker). Lane 2: DynaMarker Prestain Marker for Small RNA Plus (Diagnocine). Lane 3: ribosome footprint sample. In this study, gel slices corresponding to the 20-30-nt range (marked by the bracket) were excised. **(D)** cDNA purification using a 10% TBE-urea gel. Lane 1: ssDNA ladder. Lane 2: cDNA sample. The bracket marks the expected cDNA length (94-104 nt), and the 74-nt band corresponds to the unused RT primer. **(E)** The resulting library after amplification with 11 cycles of PCR and resolved on an 8% TBE gel. Lane 1: 20-bp ladder. Lane 2: the library product; the bracket marks the expected library size (162-172 bp). The asterisk marks the product from an unextended RT primer, which should be avoided. **(F)** Fragment Analyzer profile showing enrichment of the expected library size. LM and UM are the vendor’s internal size markers.

There are several important considerations in the major steps of ribosome footprinting: 1) Lysate preparation: it is critical to immobilize ribosomes on mRNAs to reflect their *in vivo* translational status. In eukaryotes, immobilization is typically achieved via freezing with liquid nitrogen and lysing the cells in the presence of the translation inhibitors cycloheximide (for cytosolic ribosomes) and chloramphenicol (for plastid and mitochondrial ribosomes). 2) Ribonuclease digestion: This step is the most critical for determining the nucleotide resolution of Ribo-seq. RNase I digestion may be performed without pre-purifying polysomes (Hsu et al., 2016; Chen et al., 2022; Wu and Hsu, 2022). Plant materials with fewer active ribosomes, such as adult leaves, may require prior polysome enrichment (Sotta et al., 2021). Adjusting the pH and ionic strength of the lysis buffer and titrating the amount of RNase I help maximize the nucleotide resolution (Hsu et al., 2016; Wu et al., 2019). 3) Monosome isolation: size exclusion columns, sucrose cushions, or sucrose gradients can be used to purify monosomes, but the latter two require ultracentrifugation and specialized equipment. 4) RF size selection: this has a profound effect on the Ribo-seq reads obtained and should be carefully evaluated depending on the study objective. Our previous and other studies select footprints of 28-30 nt (Hsu et al., 2016; Chen et al., 2022; Wu and Hsu, 2022), which is the major length of cytosolic RFs. Some studies isolate a wider range of footprint lengths (e.g., 20-40 nt) to capture plastid ribosomes (∼32 nt), as well as a subset of cytosolic ribosomes (21∼22 nt) (Chotewutmontri and Barkan, 2016). 5) rRNA depletion: without rRNA depletion, around 90% of the resulting Ribo-seq reads would comprise contaminant rRNA fragments (Zinshteyn et al., 2020). Two common approaches to deplete rRNAs are i) using a commercial illumina RiboZero kit, and ii) subtractive hybridization of biotinylated oligos customized to specific tissues and library preparations, which requires a pilot Ribo-seq experiment to identify the abundant contaminant sequences.

### Construction of Ribo-seq libraries using a custom method

We previously used illumina Ribo-seq kit and standalone RiboZero in our Ribo-seq protocol (Hsu et al., 2016; Wu and Hsu, 2022). The discontinuation of these reagents prompted us to test an alternative library construction method. Here, we describe a protocol that combines two custom library construction methods originally designed for yeast and bacteria (McGlincy and Ingolia, 2017; Li et al., 2021) and adapts them for plants. We applied this new protocol to 7-day-old Arabidopsis seedlings and assessed the quality of the resulting libraries. The method and a step-by-step protocol with detailed information about the reagents, as well as the analysis code, are provided in **Supplemental Files 1-4**.

Figure 1 shows the workflow of our new Ribo-seq method. We followed our optimized ribosome footprinting method, which yields strong 3-nt periodicity for open reading frame (ORF) identification (Hsu et al., 2016; Wu and Hsu, 2022). After isolating monosomes and purifying RNA, a distinct band corresponding to ∼28 nt could be observed in a polyacrylamide gel (**Figures 1C and S1**). Instead of selecting 28∼30-nt RFs, in this study, we selected 20-30-nt RFs to capture both 21∼22-nt and 28-nt RFs. Our new library construction workflow is summarized in **Figure 1B**. Briefly, the RFs are end repaired to allow their 3’ end to ligate to a pre-adenylated linker. Unligated linkers are removed by 5’ deadenylase and RecJ (an ssDNA exonuclease acting in the 5’-to-3’ direction). Following rRNA depletion by RiboZero (available as a component in the illumina TruSeq RNA-seq kit for plants), a reverse transcription (RT) primer that complements the linker enables RF cDNA synthesis. After the RNA is degraded and the cDNA is gel-purified from unutilized RT primers (**Figure 1D**), the resulting cDNA is circularized. qPCR is performed as recommended (McGlincy and Ingolia, 2017) to quantify the circularized cDNA and estimate the amount of input and number of PCR cycles needed next. Finally, library construction PCR is performed, where index and illumina sequencing sequences are incorporated. The libraries are gel-purified and evaluated with Fragment Analyzer before sequencing (**Figure 1E and 1F**).

### Evaluation of high-quality Ribo-seq data

The sequencing data were first pre-processed, and common contaminant sequences (**Supplemental File S3**) from rRNAs/tRNAs/snRNAs/snoRNAs and a few overrepresented non-coding RNAs were removed. Three technical replicates showed excellent correlations (**Figure S2**). We pooled all three replicates and used Ribo-seQC (Calviello et al., 2019) to assess data quality. As expected, for the nucleus-encoded genes, the major RF length was 28 nt (**Figure 2A**). A minor peak at 21 nt, which represents the ribosomes in the rotated conformation (Lareau et al., 2014), was also observed (**Figure 2A**). For the plastid-encoded genes, 23-27 nt RFs were enriched (**Figure 2A**). Consistent with our expectation, most of the RFs mapped to coding sequences across all RF lengths (**Figure 2B**). In the metaplot where RFs map near the start codons, the middle of coding sequence, and stop codons of protein-coding genes (**Figure 2C**), strong 3-nt periodicity (91.03%) was present within the coding sequences, and sparse reads were present within the 5’ and 3’ UTRs, as expected. Overall, our new protocol yields high-quality Ribo-seq data considering RF length distribution, genomic location distribution, and fraction of RFs enriched in the expected reading frame.

**Figure 2.**
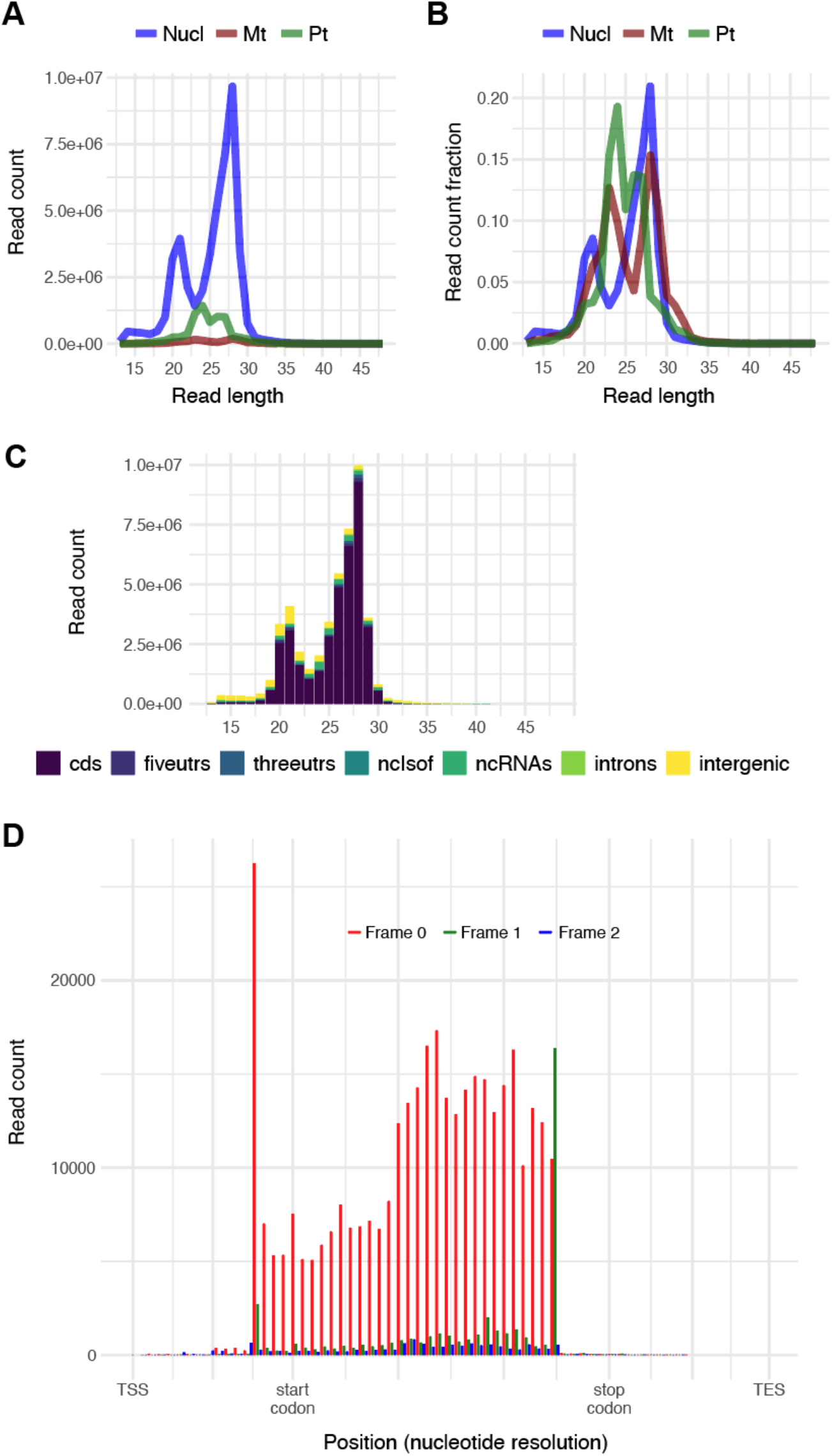
Assessments of Ribo-seq data. **(A-B)** Read length distribution of nucleus-, mitochondria- and plastid-encoded transcripts displayed based on read counts (A) or fractions (B). **(C)** Genomic features of ribosome footprints grouped by read length. Different genomic features are shown with different colors. Ribosome footprints that mapped to nucleus-encoded genes are presented. **(D)** Metaplot of protein-coding transcripts showing strong 3-nt periodicity and high enrichment within expected coding regions. Reads mapped to the three reading frames are shown in red, green and blue. The ribosome footprint position is shown by the first nucleotide of the footprint; thus, the first peak is 12 nt upstream of the start codon, consistent with our previous datasets (Hsu et al., 2016; Wu et al., 2019; Wu and Hsu, 2022).

### Considerations for future Ribo-seq experiments

Compared to our previous narrow RF size selection (28-30 nt), selecting a wider range of sizes (20-30 nt) in this study allowed us to obtain 21-nt RFs in the rotated conformation (Lareau et al., 2014). The tradeoffs are that we had more contaminating sequences, especially at 25 nt (**Figures S3 and S4**), and shorter RFs are expected to have a lower mapping rate. Depending on the purpose of the study, one might prefer to select different RF sizes. For studies focusing on cytosolic translation, one might omit chloramphenicol in the lysis buffer. If desired, one could consider including a unique molecular identifier, i.e., degenerate sequences, at the 5’ end of the linker and/or the 5’ end of the RT primer to reduce and correct bias introduced by ligation and PCR amplification (McGlincy and Ingolia, 2017; Chen et al., 2022). In summary, we have implemented a custom library construction method for our optimized ribosome footprinting protocol in Arabidopsis. This method should be readily applicable to other plant species.

## SUPPLEMENTAL DATA

The following materials are available in the online version of this article.

**Figures S1-S4**

**Supplemental File 1: Materials and methods**

**Supplemental File 2: Step-by-step Ribo-seq protocol**

**Supplemental File 3: Contaminating sequences from rRNAs, tRNAs, snRNAs, snoRNAs and dominant noncoding RNAs**

**Supplemental File 4: Code used in this study**

## AVAILABILITY OF DATA AND MATERIALS

The raw sequencing data generated in this study have been submitted to the NCBI Sequence Read Archive (SRA) under BioProject ID PRJNA854638.

## AUTHOR CONTRIBUTIONS

HLW and PYH designed the research, PYH performed the sequencing experiments, HLW analyzed the sequencing data, and HLW and PYH interpreted the results and wrote the paper.

## ACKNOWLEDGEMENTS

This work used the Vincent J. Coates Genomics Sequencing Laboratory at UC Berkeley, supported by NIH Instrumentation Grant S10 OD018174. This research was supported by a Michigan State University startup grant and an NSF award (2051885) to PYH.

