## Supplemental figures for "A custom library construction method for super-resolution ribosome profiling in Arabidopsis"

**Figure S1**

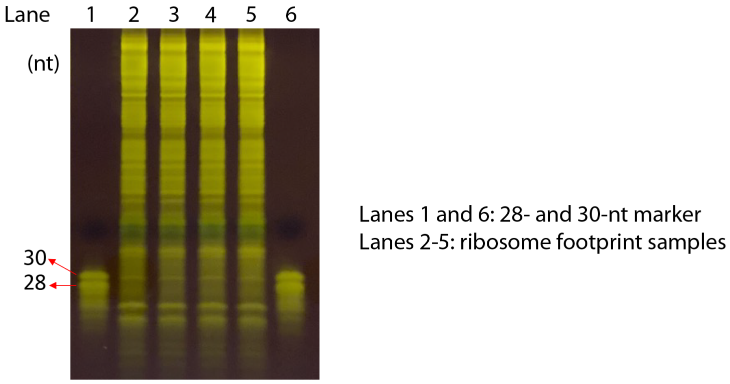


**Figure S1. Precise ribonuclease digestion in Arabidopsis yields a clear band between 28 and 30 nt.** About 200 ng of RNA (after monosome isolation) was separated on a 15% TBE-urea gel to evaluate the digestion pattern.

### Figure S2

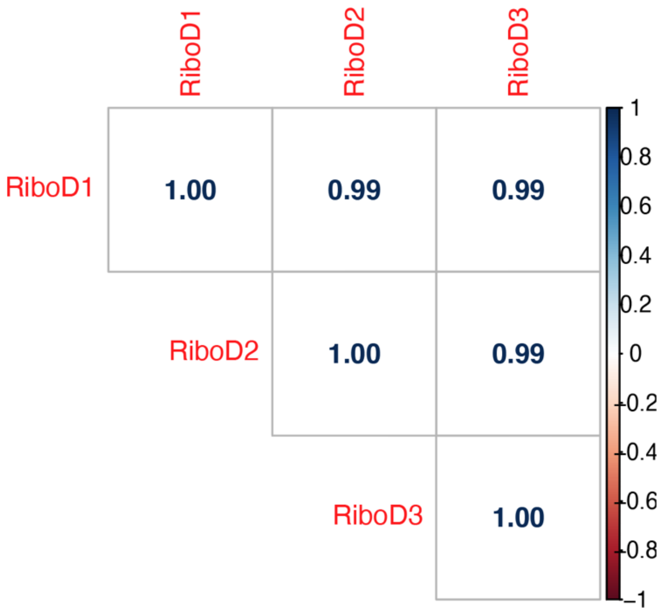


**Figure S2. High correlations among Ribo-seq data sets from three technical replicates.** Ribo-seq reads mapped to individual transcripts (considering transcripts per million, TPM) were compared among three technical replicates.

#### Figure S3

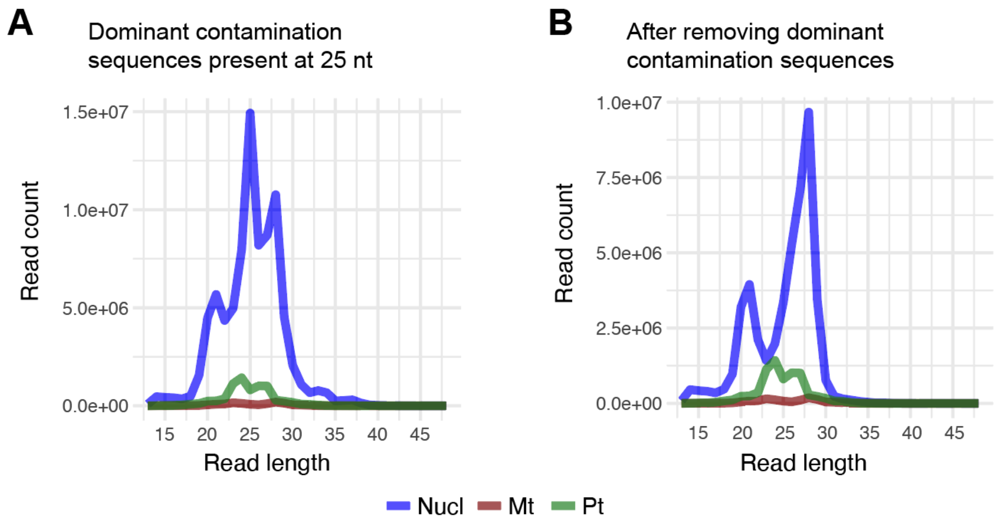


**Figure S3. Abundant contaminating sequences present at 25 nt.** (A) Read length distribution after removing rRNA, tRNA, snRNA and snoRNA contaminants. A major peak at 25 nt mainly mapped to the non-coding RNAs *AT3G06365* and *AT2G03875***.** (B) Read length distribution after removing the additional *AT3G06365* and *AT2G03875* contaminants.

**Figure S4**

| | Total reads (M) | Contam. (M) | Contam. % | Non-contam. (M) | Non-contam. % | Mapped reads (M) | Mapped % in non-contam. | Mapped % in total reads |
|---|---|---|---|---|---|---|---|---|
| Rep-1 | 49.9 | 29.8 | 59.8 | 20.1 | 40.2 | 12.7 | 63.4 | 25.5 |
| Rep-2 | 51.2 | 30.2 | 58.9 | 21.0 | 41.1 | 12.9 | 61.5 | 25.2 |
| Rep-3 | 42.0 | 23.6 | 56.2 | 18.4 | 43.8 | 11.4 | 62.1 | 27.2 |

**Figure S4. Mapping statistics from three technical replicates.** Contaminating sequences (Contam.) include rRNAs, tRNAs, snRNAs, snoRNAs and abundant non-coding RNAs (i.e., *AT3G06365* and *AT2G03875*).
