## Supplemental File S1 for "A custom library construction method for super-resolution ribosome profiling in Arabidopsis"

### SUPPLEMENTAL FILE 1: MATERIALS AND METHODS

Below, we describe the key steps in a ribosome profiling experiment and subsequent data analysis. A step-by-step Ribo-seq protocol is provided in **Supplemental File 2**. Common contaminating sequences are provided in **Supplemental File 3**. The code for the data analysis is provided in **Supplemental File 4.**

#### Plant materials

Seven-day-old Arabidopsis seedlings were grown hydroponically in sterile liquid media (2.37 g/L LS pH 5.7, 1% sucrose, 0.5 g/L MES) with shaking at 85 rpm under a 16-h light (~55 µmol m−2·s−1 from cool white fluorescent bulbs) and 8-h dark cycle at 22°C. Whole seedlings were harvested at Zeitgeber time 4 (4 h after lights on). After removing excess media with paper towels, the plant materials were immediately placed in foil, flash-frozen in liquid nitrogen, and used for lysate preparation.

#### Lysate preparation

The lysates and ribosome footprints were prepared according to our previous methods (Hsu et al., 2016; Wu and Hsu, 2022). Ice-cold lysis buffer (100 mM Tris-HCl [pH 8], 40 mM KCl, 20 mM MgCl2, 2% [v/v] polyoxyethylene [10] tridecyl ether [Sigma, P2393], 1% [w/v] sodium deoxycholate [Sigma, D6750], 1 mM dithiothreitol, 100 µg/mL cycloheximide [Sigma, C4859], 100 µg/mL chloramphenicol [Sigma R4408], and 10 units/mL DNase I [Lucigen, D9905K]) was prepared and aliquoted in 5-mL centrifuge tubes. After grinding the frozen tissue to a fine powder with a mortar and a pestle, the powder was swept into the aliquoted lysis buffer (0.1 g per 400 µL lysis buffer) and immediately resuspended by vortexing. The lysates were mixed at 4°C for 10 minutes with shaking, then spun at 5,000 × g for 3 min at 4°C. The supernatant was transferred to new tubes and spun at 20,000 × g for 10 min at 4°C. The supernatant was transferred to new tubes again, and 200-µL (for ribosome footprint preparation) and 50-µL (for total RNA extraction) aliquots were made. The aliquots were frozen in liquid nitrogen and saved at -80°C until further processing.

**Ribosome footprinting, monosome isolation and size selection**

The 200-μL lysate aliquots above were processed to generate ribosome footprints. The RNA concentration was determined via Qubit RNA high-sensitivity assay (Thermo Fisher Scientific, Q32855) using 10-fold diluted lysate. RNase I (Lucigen N6901K, 50 U/per 40 μg RNA) was added to the lysates, and the reactions were mixed on a nutator at room temperature for 1 hour. The reactions were terminated by placing the samples on ice and adding 15 μL SUPERase-In (Thermo Fisher Scientific AM2696). Monosomes were isolated by applying each 100 μL of digested lysate onto one size exclusion column (illustra MicoSpin S-400 HR, GE Healthcare 27-5140-01), which was equilibrated with 3 mL of polysome buffer (100 mM Tris-HCl [pH 8], 40 mM KCl, 20 mM $MgCl_2$) in advance. Then, 10 μL of 10% SDS was added, and RNA >17 nt was isolated using an RNA Clean & Concentrator kit (Zymo Research R1015). The purified RNA was separated via 15% (w/v) TBE-urea PAGE (Thermo Fisher Scientific; EC68852BOX), and gel slices corresponding to 20-30 nt were excised. Ribosome footprints were recovered and used for library construction.

**Ribo-seq library construction**

The library construction method was modified from two methods (McGlincy and Ingolia, 2017; Li et al., 2021). The ribosome footprints were repaired via T4 PNK (NEB M0201S) in the absence of ATP and ligated to a universal miRNA cloning linker (NEB S1315S) with T4 RNA Ligase 2 truncated K227Q (NEB M0351S). The excess linkers were removed with 5´ deadenylase (NEB M0331S) and RecJf (NEB M0264S). After purifying the ligation products using an Oligo Clean & Concentrator kit (Zymo D4061), rRNA depletion was performed using RiboZero, which is included in the TruSeq Stranded Total RNA Library Prep Plant kit (illumina 20020610), and the ligated ribosome footprints were purified again with an Oligo Clean & Concentrator kit. Next, reverse transcription was carried out with ProtoScript II (NEB M0368L) and a reverse transcription primer whose sequence at the 3' end was complementary to the linker. The RNA was degraded by treating the sample with NaOH at a final concentration of ~0.1M. The cDNA was purified again with an Oligo Clean & Concentrator kit and separated via 10% (w/v) TBE-urea PAGE (Thermo Fisher Scientific EC68752BOX). Then, cDNA of the expected size (94-104 nt) was selected, recovered, and circularized using CircLigase (Lucigen CL4111K).

**qPCR quantification of circularized cDNA**

The circularized cDNA was quantified via qPCR using Luna universal qPCR master mix (NEB M3003S). A synthesized oligo was used as a positive control and to establish the standard curve for quantification. The primer and oligo sequences are listed in **Supplemental File 2**.

**Library PCR amplification, purification, and sequencing**

Eleven-cycle library PCR with indexed primers was performed using Phusion high-fidelity PCR master mix (NEB M0531S) following the recommendations based on the qPCR quantification (McGlincy and Ingolia, 2017). The PCR products were separated on an 8% TBE gel (Thermo Fisher Scientific EC62152BOX), and the library of the expected size (162-172 bp) was selected and recovered. The size of the recovered libraries was evaluated using Fragment Analyzer (Agilent). The libraries were quantified via Qubit dsDNA HS assay (Thermo Fisher Scientific Q32854) and pooled at equal molarity. Single-end 50-bp sequencing was performed in a HiSeq 4000. The raw sequencing data have been deposited in the NCBI Sequence Read Archive (SRA) under BioProject ID PRJNA854638.

**Data analysis** (The code used in this study is provided in **Supplemental File 4.**)

**Step 1 (Trim adaptors and remove low-quality sequences)**: The adaptor sequence (CTGTAGGCACCATCAAT) was first trimmed from the Ribo-seq reads with *fastx_clipper* (*fastx toolkit* v0.11.7, http://hannonlab.cshl.edu/fastx_toolkit/). For the *fastx_clipper* function, the -c, -n, -v, -Q33 options were used to discard unclipped reads, keep reads with unknown nucleotides, and provide verbose output, and the Q33 filter was used to remove low-quality reads.

**Step 2 (Remove contaminating sequences with *Bowtie2*):** We first built a *Bowtie2* (v. 2.3.4.1) (Langmead and Salzberg, 2012) index with the *bowtie2-build* function for removing unwanted contaminating sequences such as those from rRNAs, tRNAs, snRNAs, and snoRNAs. We also used the same method to remove additional high-abundance non-coding RNAs (i.e., *AT3G06365* and *AT2G03875*). These contaminating sequences are listed in **Supplemental File 3**. We next used *Bowtie2* with seed length (-L option) 20 to extract Ribo-seq sequences that did not map to contaminating sequences.

**Step 3 (Map Ribo-seq reads):** We then created an index file for *STAR* aligner (v 6.2.0c) (Dobin et al., 2013) against the Araport11 transcriptome and the TAIR10 genome using the following options: --runMode genomeGenerate, --sjdbOverhang 34.

We mapped the remaining Ribo-seq reads from Step 2 with *STAR* aligner using the following options: --alignIntronMax 5000, --alignIntronMin 15, --outFilterMismatchNmax 1, --outFilterMultimapNmax 20, --outFilterType BySJout, --alignSJoverhangMin 8, --alignSJDBoverhangMin 2, --outSAMtype BAM SortedByCoordinate, --quantMode TranscriptomeSAM, --outSAMmultNmax 1, --outMultimapperOrder Random.

**Step 4 (Conduct Ribo-seQC analysis):** Next, we used the Ribo-seQC package (Calviello et al., 2019) to analyze the quality of the Ribo-seq reads. We first used the *export.2bit* function from the *rtracklayer* package (Lawrence et al., 2009) to create the 2bit file for the TAIR10 genome. We generated the annotation file for *Ribo-seQC* with the *prepare_annotation_files* function using the Araport11 gtf file and the genome 2bit file as inputs. The *Ribo-seQC* output was generated using the *RiboseQC_analysis* function with the *Ribo-seQC* annotation file and the bam file generated in Step 3.

**Step 5 (Conduct Kallisto quantification for Ribo-seq data and correlation analysis)**: We used Kallisto (Bray et al., 2016) to create the index and quantify the three technical replicates. We then used the corrplot function from the corrplot library in R (v4.0.3) (R Core Team (2013), 2017) to plot the correlation of the three replicates.

**Step 6 (Calculate 3-nt periodicity)**: We used the output file (ending with bam_results_RiboseQC) from Ribo-seQC to calculate 3-nt periodicity in R. A total of 93 nucleotides (31 codons) were considered, including 33 nucleotides starting from the start codon, 33 nucleotides in the middle of the transcript, and 27 nucleotides from -2 to -10 codons upstream of the stop codon.
