## Supplemental File S2 for "A custom library construction method for super-resolution ribosome profiling in Arabidopsis"

**SUPPLEMENTAL FILE 2: Step-by-step Ribo-seq protocol**

**REAGENTS**

| Material | Manufacturer | Catalog No. |
|---|---|---|
| RNaseZap RNase Decontamination Solution (Optional) | Thermo Fisher Scientific | AM9780 |
| Liquid nitrogen | | |
| Mortar, 50 mL, 47 x 70 mm | VWR | 89038-144 |
| Pestle, 50 mL, 114 mm | VWR | 89038-160 |
| Nuclease-Free Water (not DEPC-Treated) | Thermo Fisher Scientific | 4387936 |
| Sodium deoxycholate | Sigma | D6750-100G |
| Polyoxyethylene (10) tridecyl ether | Sigma | P2393-100G |
| 2 M KCl | Thermo Fisher Scientific | AM9640G |
| 1 M $MgCl_2$ | Thermo Fisher Scientific | AM9530G |
| 1 M Tris pH 8.0 | Thermo Fisher Scientific | AM9855G |
| 10X TBE Buffer | Thermo Fisher Scientific | AM9863 |
| Chloramphenicol, ready made solution | Sigma | R4408-10ML |
| Cycloheximide, ready made solution | Sigma | C4859-1ML |
| DL-dithiothreitol (DTT) solution, BioUltra | Sigma | 43816-10ML |
| RNase-free DNase I | Lucigen | D9905K |
| RNase I, E. coli | Lucigen | N6901K |
| SUPERase-In | Thermo Fisher Scientific | AM2696 |
| illustra MicroSpin S-400 HR Columns | GE Healthcare | 27-5140-01 |
| 10% SDS solution | Thermo Fisher Scientific | AM9822 |
| RNA Clean & Concentrator-5 no DNase I | Zymo | R1015 |
| Oligo Clean & Concentrator | Zymo | D4061 |

| | | |
|---|---|---|
| DNA Clean & Concentrator-5 (Capped) | Zymo | D4013 |
| 5 M NaCl | Thermo Fisher Scientific | AM9760G |
| 3M NaOAc pH 5.5 | Thermo Fisher Scientific | AM9740 |
| 0.5 M EDTA pH 8.0 | Thermo Fisher Scientific | AM9260G |
| Ethanol, 200 proof | Decon Labs | V1016 |
| Isopropanol, molecular biology grade | Fisher | BP2618500 |
| GlycoBlue (15 mg/mL) | Thermo Fisher | AM9515 |
| 15% TBE-urea gel 12 well | Thermo Fisher Scientific | EC68852BOX |
| 10% TBE-urea gel 12 well | Thermo Fisher Scientific | EC68752BOX |
| 8% TBE gel, 12 wells | Thermo Fisher Scientific | EC62152BOX |
| Disposable transfer pipette | Fisher | 13-711-20 |
| DynaMarker, Prestain Marker for Small RNA Plus | Diagnocine | DM253 |
| miRNA marker (optional) | NEB | N2102S |
| 20/100 Ladder | IDT | 51-05-15-02 |
| 20 bp Ladder | Bayou Biolabs | L-100 |
| Gel loading buffer II (Denaturing PAGE) | Thermo Fisher Scientific | AM8546G |
| Gel loading dye, purple (6X) | NEB | B7024S |
| SYBR Gold nucleic acid gel stain 500 µL | Thermo Fisher Scientific | S11494 |
| TruSeq Stranded Total RNA Library Prep Plant | illumina | 20020610 |
| QUBIT RNA HS assay kit | Thermo Fisher Scientific | Q32855 |
| QUBIT dsDNA HS assay | Thermo Fisher Scientific | Q32854 |
| T4 polynucleotide kinase (PNK) | NEB | M0201S |

| | | |
|---|---|---|
| Universal miRNA cloning linker | NEB | S1315S |
| T4 RNA Ligase 2 truncated K227Q | NEB | M0351S |
| 5´ Deadenylase | NEB | M0331S |
| RecJf | NEB | M0264S |
| ProtoScript II | NEB | M0368L |
| dNTP (10mM each) | Thermo Fisher Scientific | R0191 |
| Sodium hydroxide | Fisher | S318500 |
| CircLigase | Lucigen | CL4111K |
| Luna universal qPCR master mix | NEB | M3003S |
| Phusion high-fidelity PCR master mix with HF buffer | NEB | M0531S |
| Olympus 5ml Centrifuge tube | Genesee | 24-285S |
| Ultrafree-MC GV centrifugal filter | EMD Millipore | UFC30GV00 |
| MAXYMum Recovery PCR tubes | VWR | 22234_056 |
| Non-sticky RNase-free tubes 1.5 mL | Thermo Fisher Scientific | 50591363 |
| 10µL pipette filter tips, low binding, sterile | Genesee | 24_401 |
| 20µL pipette filter tips, low binding, sterile | Genesee | 24_404 |
| 200µL pipette filter tips, low binding, sterile | Genesee | 24_412 |
| 1000µL pipette filter tips, low binding, sterile | Genesee | 24_430 |
| 200 µL round gel tip, 0.58 mm | Genesee | 14_101 |
| MicroAmp fast optical 96-well reaction plate | Thermo Fisher Scientific | 4346906 |
| TempPlate RT qPCR sealing film | USA Scientific | 2978-2100 |
| 0.2 mL PCR 8-tube strip with 8-cap strips | Genesee | 24_705 |

### EQUIPMENT

| Equipment | Manufacturer | Catalog No. |
|---|---|---|
| Refrigerated centrifuge for 5 mL tubes and 1.5 mL tubes | Eppendorf | 5430R |
| BD Clay Adams Nutator Mixer | VWR | 15172-203 |
| Qubit fluorometer | Thermo Fisher Scientific | Q33238 |
| Mini gel tank | Thermo Fisher Scientific | A25977 |
| Electrophoresis power supply | Fisher | FB300Q |
| DarkReader (Dark Field Transilluminator) | Clare Chemical Research | DR46B |
| T100 thermocycler | Bio-Rad | 1861096 |
| Magnetic Separation Stand, 12 position, 1.5 mL | Promega | Z5342 |
| QuantStudio 3 Real-Time PCR System | Thermo Fisher Scientific | A28567 |

### OLIGOS (all except the Universal miRNA cloning linker can be ordered from IDT)

- All oligos should be dissolved in nuclease-free water (NF-H2O) at the appropriate concentrations, aliquoted, and stored at -20°C.
- The linker sequence is shown in red, and the sequence complementary to the linker is shown in red and underlined.
- The forward library PCR primer sequence is shown in blue
- The shared sequence in the reverse qPCR primer and the indexed reversed library PCR primer is shown in green; its complementary sequence is shown in green and underlined

| Oligo name | Scale & Purification | Sequence | Conc. |
|---|---|---|---|
| Universal miRNA cloning linker (5′ adenylated, 3′ blocked) | NEB S1513S | (5´) rAppCTGTAGGCACCATCAAT-NH2 (3´) | 20 μM |

| | | | |
|---|---|---|---|
| Reverse transcription primer | 100 nm, HPLC | /5Phos/AGATCGGAAGAGCGTCGTGTAGGGAAAGAGTGT/iSp18/CAAGCAGAAGACGGCATACGAGATATTGATGGTGCCTACAG | 1.25 µM |
| Forward library PCR primer | 25 nm, standard desalt | 5'-CAAGCAGAAGACGGCATACGA-3' | 10 µM |
| Reverse qPCR primer | 25 nm, standard desalt | 5'-ACACTCTTTCCCTACACGACG-3' | 10 µM |
| Positive control for qPCR (100 nt including **26-nt-synthetic/hypothetical RF sequence)** | 4 nm ultramer, standard desalt | 5'CAAGCAGAAGACGGCATACGAGATATTGATGGTGCCTACAG**TCGCATTACCCTGTTATCCCTAACAT**AGATCGGAAGAGCGTCGTGTAGGGAAAGAGTGT3' | 10 µM |
| Indexed reverse library PCR primer 1 | 4 nm ultramer, standard desalt | 5'AATGATACGGCGACCACCGAGATCTACACGATCGGAAGAGCACACGTCTGAACTCCAGTCAC**ATCACG**ACACTCTTTCCCTACAC3' | 10 µM |
| Indexed reverse library PCR primer 2 | 4 nm ultramer, standard desalt | 5'AATGATACGGCGACCACCGAGATCTACACGATCGGAAGAGCACACGTCTGAACTCCAGTCAC**CGATGT**ACACTCTTTCCCTACAC3' | 10 µM |
| Indexed reverse library PCR primer 3 | 4 nm ultramer, standard desalt | 5'AATGATACGGCGACCACCGAGATCTACACGATCGGAAGAGCACACGTCTGAACTCCAGTCAC**TTAGGC**ACACTCTTTCCCTACAC3' | 10 µM |
| Indexed reverse library PCR primer 4 | 4 nm ultramer, standard desalt | 5'AATGATACGGCGACCACCGAGATCTACACGATCGGAAGAGCACACGTCTGAACTCCAGTCAC**TGACCA**ACACTCTTTCCCTACAC3' | 10 µM |
| Indexed reverse library PCR primer 5 | 4 nm ultramer, standard desalt | 5'AATGATACGGCGACCACCGAGATCTACACGATCGGAAGAGCACACGTCTGAACTCCAGTCAC**ACAGTG**ACACTCTTTCCCTACAC3' | 10 µM |

| | | | |
|---|---|---|---|
| Indexed reverse library PCR primer 6 | 4 nm ultramer, standard desalt | 5'AATGATACGGCGACCACCGAGATCTACACGATCGGAAGAGCACACGTCTGAACTCCAGTCAC**GCCAAT**ACACTCTTTCCCTACAC3' | 10 µM |
| Indexed reverse library PCR primer 7 | 4 nm ultramer, standard desalt | 5'AATGATACGGCGACCACCGAGATCTACACGATCGGAAGAGCACACGTCTGAACTCCAGTCAC**CAGATC**ACACTCTTTCCCTACAC3' | 10 µM |
| Indexed reverse library PCR primer 8 | 4 nm ultramer, standard desalt | 5'AATGATACGGCGACCACCGAGATCTACACGATCGGAAGAGCACACGTCTGAACTCCAGTCAC**ACTTGA**ACACTCTTTCCCTACAC3' | 10 µM |
| Indexed reverse library PCR primer 9 | 4 nm ultramer, standard desalt | 5'AATGATACGGCGACCACCGAGATCTACACGATCGGAAGAGCACACGTCTGAACTCCAGTCAC**GATCAG**ACACTCTTTCCCTACAC3' | 10 µM |
| Indexed reverse library PCR primer 10 | 4 nm ultramer, standard desalt | 5'AATGATACGGCGACCACCGAGATCTACACGATCGGAAGAGCACACGTCTGAACTCCAGTCAC**TAGCTT**ACACTCTTTCCCTACAC3' | 10 µM |
| Indexed reverse library PCR primer 11 | 4 nm ultramer, standard desalt | 5'AATGATACGGCGACCACCGAGATCTACACGATCGGAAGAGCACACGTCTGAACTCCAGTCAC**GGCTAC**ACACTCTTTCCCTACAC3' | 10 µM |
| Indexed reverse library PCR primer 12 | 4 nm ultramer, standard desalt | 5'AATGATACGGCGACCACCGAGATCTACACGATCGGAAGAGCACACGTCTGAACTCCAGTCAC**CTTGTA**ACACTCTTTCCCTACAC3' | 10 µM |

**REAGENT SETUP:**

- All reagents and supplies should be handled with care to ensure an RNase-free environment.
- Nuclease-free water is abbreviated "NF-H2O"

1. Cycloheximide stock solution (100 mg/mL), aliquot and store at -80°C
2. Chloramphenicol stock solution (100 mg/mL), aliquot and store at -80°C
3. 20% (v/v) PTE (polyoxyethylene (10) tridecyl ether): dispense 10 mL in a 50-mL sterile centrifuge tube, dissolve in NF-H2O and adjust to a final volume of 50 mL, store at room temperature
4. 10% (w/v) sodium deoxycholate: weigh 5 g in a sterile 50-mL centrifuge tube, dissolve in NF-H2O and adjust to a final volume of 50 mL, store at room temperature
5. 1 M DTT, aliquot and store at -80°C
6. 1 M NaOH: dissolve 0.04 g in 1 mL NF-H2O, store at room temperature
7. RNA extraction buffer, store at room temperature

| | 50 mL | Final concentration |
|---|---|---|
| NF-H2O | 43.65 | - |
| 3M NaOAc, pH5.5 | 5 | 300 mM |
| 10% SDS | 1.25 | 0.25% |
| 0.5M EDTA | 0.1 | 1 mM |

8. DNA extraction buffer, store at room temperature

| | 50 mL | Final concentration |
|---|---|---|
| NF-H2O | 46.4 | - |
| 5M NaCl | 3 | 300 mM |
| 1M Tris, pH 8 | 0.5 | 10 mM |
| 0.5M EDTA | 0.1 | 1 mM |

9. NEB Universal miRNA Cloning Linker:
    - Resuspend in 43 µL NF-H2O (final 20 µM)
    - Aliquot and store at -20°C
10. Tris (10 mM, pH8): dilute 1 M Tris pH 8 stock with NF-H2O, store at room temperature

**PROCEDURES:**

- All heating steps are performed in a thermocycler
- Starting with step 27, use low-bind (or non-sticky) 1.5-mL tubes and PCR tubes

**A. Preparation of plant lysates**

1. Make lysis buffer (add cycloheximide, chloramphenicol, DNase I, and DDT fresh); chill the buffer on ice:

| | 1 mL | | 50 mL | | Final conc. |
|---|---|---|---|---|---|
| Nuclease-free water (NF-H2O) | 647 | μL | 32.35 | mL | - |
| 1M Tris-HCl, pH 8 | 100 | μL | 5 | mL | 100 mM |
| 20% Polyoxyethylene (10) tridecyl ether **(before Sodium deoxycholate)** | 100 | μL | 5 | mL | 2% |
| 10% Sodium deoxycholate | 100 | μL | 5 | mL | 1% |
| 2M KCl | 20 | μL | 1 | mL | 40 mM |
| 1M MgCl2 | 20 | μL | 1 | mL | 20 mM |
| 1M DTT | 1 | μL | 50 | mL | 1 mM |
| 100 mg/mL Cycloheximide | 1 | μL | 50 | mL | 100 μg/mL |
| 100 mg/mL Chloramphenicol | 1 | μL | 50 | mL | 100 μg/mL |
| DNase I (1U/μL) | 10 | μL | 500 | mL | 10 U/mL |

2. Aliquot the lysis buffer into 5 mL centrifuge tubes: 400 μL lysis buffer is needed for 0.1 g of whole-seedling Arabidopsis samples.
3. Collect samples in aluminum foil and freeze in liquid nitrogen immediately.
4. (Chill the centrifuges) Tare the 5 mL tube with lysis buffer on a scale first. Then, grind the samples using a chilled mortar and pestle with liquid nitrogen. Sweep the ground tissue into the buffer and weigh. Work quickly to prevent the tissue from thawing. Add more lysis buffer if needed to keep the tissue: buffer ratio consistent. Vortex to thoroughly resuspend the tissue. Leave the sample on ice while processing the other samples.
5. Vortex all samples again. Shake the samples at 4°C (on ice or in a cold room) for 10 min.
6. Spin shoot samples at 5,000 × g at 4°C for 3 min. During the spin, prepare new 1.5-mL tubes for the next step and make holes in the ice with a spare 5-mL tube to avoid disturbing the tissue debris after the spin.

7. Transfer the supernatant to chilled 1.5-mL tubes. Centrifuge at 20,000 × g at 4°C for 10 min. During the spin, prepare new 1.5-mL tubes for the next step and make holes in the ice with a spare 1.5-mL tube to avoid disturbing the tissue debris after the spin.
8. Transfer the supernatant to new chilled 1.5-mL tubes.
9. (Optional) Quantify the RNA concentration with a Qubit RNA HS assay using 10X diluted lysate.
10. (Optional) Adjust the samples to the same RNA concentration using lysis buffer.
11. Make aliquots of 200 µL (for ribosome footprint samples) and 50 µL (for RNA samples). Flash freeze the aliquots with liquid nitrogen and store at -80°C.

### B. RNase I digestion & isolation of ribosome-protected fragments

12. (Do this during one of the buffer steps below if not done already) Quantify RNA concentration with a Qubit RNA HS assay using 10X diluted lysate.
13. Prepare size exclusion columns (SECs) and SEC buffer.
    - Vortex and invert the SECs several times to resuspend the resin. Remove both ends of the columns and place them on a rack to allow gravity flow.
    - Prepare 3 mL SEC buffer for each column; 1 column is used for 100 µL of lysate.

| | 1 mL | | 50 mL | |
|---|---|---|---|---|
| NF-H2O | 860 | 860 | 43 | mL |
| 1M Tris-HCl, pH 8 | 100 | 100 | 5 | mL |
| 2M KCl | 20 | 20 | 1 | mL |
| 1M MgCl2 | 20 | 20 | 1 | mL |

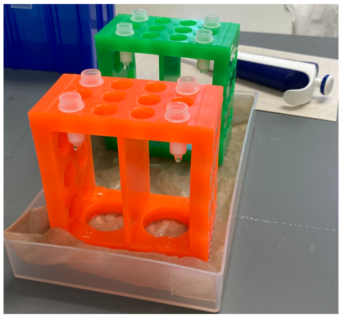
Set up SECs on a rack.

    - Resuspend the resin on the column cap using 0.5 mL SEC buffer by pipetting and adding it to the column. Initiate the flow with a gloved finger if necessary.
    - Add 0.5 mL SEC buffer every 15-20 min; after adding the buffer 3 times, set up the digestion (i.e., start step 14 right after adding the buffer for the 4th time).
    - After equilibrating the column with 3 mL of SEC buffer, tight the screw cap on all the way and then turn it ¼ turn to loosen the cap. Spin at 600 × g for 4 min before the digestion is almost done.
14. Add 50 units of RNase I per 40 µg RNA for each sample (200 µL lysate). Briefly vortex. Gently mix on a nutator for 1 hour.

15. Quickly move the samples to ice. Add 15 µL (per 200 µL lysate) of SUPERase-IN and mix.
16. Isolate monosomes by loading ~107 µL of digested lysate onto the center of the column. Spin at 600 × g for 2 min.
17. Add 10 µL of 10% SDS to each SEC elution.
18. Purify RNA > 17 nt with a modified Zymo RNA Clean & Concentrator-5 protocol
    - In step 1 of the Zymo protocol, use 290 µL RNA Binding Buffer
    - In step 2 of the Zymo protocol, use 655 µL EtOH
    - Repeat Zymo protocol step 3 to load the same sample if > 800 µL (we typically use 1 Zymo column to combine 2 SEC elutes of the same sample together; the RNA yield is within the binding capacity of the Zymo column 10 µg)
    - Continue with the purification according to the manufacturer's instructions
    - Elute with 11 µL 10 mM Tris pH 8
    - Combine 1 µL elution with 9 µL 10 mM Tris pH 8. This will be used for Qubit quantification and to check the digestion
    - The other ~10 µL elution will be used in Step 23.
19. Quantify the RNA concentration (use the 10X diluted sample above) using a Qubit RNA HS.
20. (Optional) Run 200 ng RNA (use the 10X diluted sample above) in a 15% TBE-urea gel to check the digestion
    - Prerun a 15% TBE-urea gel for 15 min
    - Prepare 450 mL 1x TBE, save ~40 mL on ice for staining
    - Mix RNA with 2x gel loading buffer II; also prepare the 28/30-nt marker (from the discontinued illumina Ribo-seq kit or custom synthesized)
    - Denature at 80°C for 90 s, put on ice immediately
    - Rinse the wells using a clean transfer pipette before loading samples
    - Run the gel at 200V for 65 min
    - Stain the gel with SYBR-GOLD in ice cold 1x TBE for 3 min (4 µL SYBR-GOLD in 40 mL TBE buffer above)
    - Image using a Dark Field Transilluminator or a UV Transilluminator
    - In our experience, a clear band between 28 and 30 nt suggests good digestion for Arabidopsis

**Evaluation of digestion patterns**

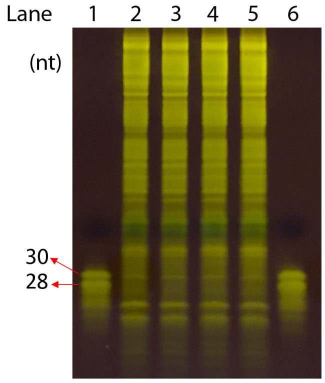


Lanes 1, 6: 28 and 30-nt marker.
Lanes 2-5: ribosome footprint samples

**C. Size selection of ribosome footprints (RFs)**

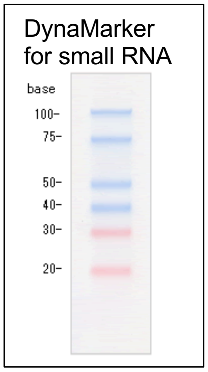


21. Prerun a 15% TBE-urea gel for 15 min
    - Prepare 400 mL 1x TBE buffer
    - Rinse the wells using a clean transfer pipette.
22. To select 20-30 nt, prepare the sRNA DynaMarker (do not heat).
23. Mix ~10 µL of RF sample from Step 18 with 10 µL of 2x gel loading buffer II.
24. Denature at 80°C for 90 s. Put on ice immediately.
25. Rinse the gel wells using a clean transfer pipette before loading the samples.
26. Separate each sample (2 wells per sample, 10 µL each) with the sRNA DynaMarker. Run at 200 V for 60 min (prepare the tubes while the gel is running).
27. Open the gel cassette and leave the gel on one side of the cassette. Put the gel and the cassette on top of a piece of white paper. Excise the gel between 20-30 nt (cut 2 wells of the same sample together) using a clean blade and place the gel slice into 1.5 mL low-bind tube.
28. Add 600 µL RNA gel extraction buffer to each sample. Make sure the gel is submerged.
29. Leave on dry ice for 30 min.
30. Thaw the samples with shaking on a nutator at room temperature overnight.
31. Briefly spin and transfer the liquid to a Ultrafree-MC GV centrifugal filter tube. Centrifuge at 2300 × g for 3 min.
32. Transfer the elution to a new 1.5-mL low-bind tube.
33. Add 2 µL of GlycoBlue and mix well.
34. Add 800 µL of isopropanol and mix well.
35. Leave the samples on dry ice for 1 hour or at -80°C overnight.
36. Centrifuge at 20,000 × g at 4°C for 30 min to pellet the RFs. (Prepare fresh 80% EtOH and leave on ice during the spin.)
37. Remove all liquid (using a 1-mL pipette first, then using a 10-µL pipette again; **DON'T spin here**).
38. Wash the pellet with 800 µL ice-cold 80% EtOH and remove the liquid without changing tips.
39. Briefly spin and remove all liquid (using a 10-µL pipette).
40. Put the tubes sideways in a microfuge tube rack. Air dry for 10 min.
41. Resuspend the RNA pellet in 3.5 µL of 10 mM Tris, pH 8.
    (Optional stopping point at -80°C)

**D. Dephosphorylation and linker ligation**

42. Prepare T4 PNK end-repair master mix on ice:

| | Volume (µL) |
|---|---|
| T4 PNK buffer w/o ATP (10X) | 0.5 |
| T4 PNK (10 U/µL) | 0.5 |
| SUPERase-IN (20 U/µL) | 0.5 |

43. Add 1.5 µL end-repair master mix to the 3.5 µL RF and mix.
44. Incubate at 37°C for 1 hr.
    (Optional stopping point at -80°C)
45. Prepare the linker ligation master mix on ice:

| | Volume (µL) |
|---|---|
| 50% w/v PEG-8000 | 3.5 |
| 10X T4 RNA ligase buffer | 0.5 |
| Universal miRNA cloning linker (20 µM) | 0.5 |
| T4 Rnl2(tr) K227Q (200 U/µL) | 0.5 |

46. Add 5 µL linker ligation master mix to the 5-µL sample above and mix by pipetting.
47. Incubate at 22°C for 3 hr in a thermocycler (set the lid temperature off).
48. Deplete unligated linkers by adding:
- 0.5 µL 5'-deadenylase (10 U/µL)
- 0.5 µL RecJ exonuclease (10 U/µL)
- 30°C for 45 min
49. Purify ligations with a Zymo Oligo Clean & Concentrator kit:
- Add 39 µL of NF-H2O to each sample; now the sample volume is 50 µL.
- Proceed with the rest of the protocol.
50. Elute in 11 µL **NF-H2O**.
51. Transfer 10 µL of sample to a low-bind PCR tube.
    (Optional stopping point at -80°C)

**E. Ribosomal RNA depletion**

52. Thaw the following reagents for RiboZero (a component in the TruSeq Stranded Total RNA Library Prep Plant kit):

| Item | Storage | Instruction |
|---|---|---|
| RRM-P (rRNA Removal Mix - Plant) | -80°C | Thaw, mix and put on ice |
| RBB (rRNA Binding Buffer) | -80°C | Thaw, mix and put on ice |
| RRB (rRNA Removal Beads) | 4°C | **Room temperature for 30 min** |

53. Save the following **RNA Denaturation program** in a thermocycler.
    - Preheat lid and set to 100°C
    - 68°C for 5 min
    - Hold at 4°C

54. Combine the following in a low-bind PCR tube

| | 1x reaction (µL) |
|---|---|
| RF from Step 51 | 10 |
| RBB (buffer) | 5 |
| RPM-P | 5 |
| Total = | 20 |

Pipet up and down 10x.

55. In a thermocycler, run the RNA Denaturation program (**prepare the RRB for Steps 57-58**).
56. Incubate at room temperature for 1 min.
57. Vortex RRB (beads) until well dispersed.
58. Aliquot 35 µL of RRB into each new 1.5-mL low-bind tubes.
59. Transfer the RF sample from Step 56 to the aliquoted RRB, pipette up and down 10x.
60. Incubate at room temperature for 1 min.
61. Place the reaction on a magnetic stand. Wait for 1 minute until the solution is clear.
62. Transfer the suspension (using 10-µL tips) to a new 1.5 mL low-bind tube. Expect to recover ~40 µL.
63. Purify RFs with a Zymo Oligo Clean & Concentrator kit:
- Add NF-H2O to each sample to a final volume of 50 µL.
- Proceed with the rest of the protocol.

- Elute in 11 µL 10 mM Tris pH 8. Transfer 10 µL to low-bind PCR tubes and proceed to the next step.

**F. Reverse transcription and cDNA purification**

64. Add 2 µL RT primer (1.25 µM) to all samples. Mix.
65. Denature at 65°C for 5 min. Put on ice immediately.
66. Set the thermocycler to 50°C.
67. Prepare the RT master mix on ice:

| | Volume (µL) | Final |
|---|---|---|
| 5X Protoscript II buffer | 4 | 1x |
| dNTPs (**10 mM each**) | 1 | 0.5 mM each |
| 0.1M DTT | 1 | 5 mM |
| SUPERase-IN (20 U/µL) | 1 | 1 U/µL |
| Protoscript II (200 U/µL) | 1 | 10 U/µL |

68. Add 8 µL RT master mix to each sample and mix. Now the volume is 20 µL.
69. Incubate at 50°C for 30 min.
70. Add 2.2 µL of **1 M NaOH** to each reaction and mix; incubate at 70°C for 20 min.
71. Purify cDNA with a Zymo Oligo Clean & Concentrator kit:
- Add NF-H2O to each sample to a final volume of 50 µL.
- Proceed with the rest of the protocol.
- Elute in 6.5 µL 10 mM Tris pH8. Transfer to low-bind PCR tubes.
72. Prerun a 10% TBE-urea gel at 200 V for 15 min in 1x TBE.
    - Prepare 450 mL 1x TBE buffer.
    - **Save 40 mL of 1x TBE buffer and leave on ice for gel staining.**
    - Rinse the wells using a clean transfer pipette.
73. Add 6.5 µL of 2x gel loading buffer II to each sample. Also prepare the 20/100 ssDNA ladder needed to separate each sample.
74. Denature the samples and ladder at 80°C for 90 s. Move to ice immediately.
75. Thoroughly rinse the gel wells, then load the samples and ladder onto a denaturing 10% TBE-urea gel.
76. Run at 200 V for 80 min in 1× TBE buffer (**put the 40 mL 1x TBE on ice**).

77. Stain the gel with SYBR-GOLD in ice-cold 1x TBE for 3 min (4 µL SYBR-GOLD in 40 mL TBE buffer above).
78. Visualize gel using a Dark Field Transilluminator. Excise the RT products between 94 and 104 nt using a clean blade (do not cut a wider range; it will increase unwanted PCR products). Place the excised gel in a 1.5-mL low-bind tube.
79. Add 500 µL **DNA** extraction buffer. Extract the cDNA as described in Steps 28-39.
80. Resuspend the cDNA in 15 µL 10 mM Tris pH 8 and transfer to a low-bind PCR tube.
    (Optional stopping point at -20°C overnight or -80°C for longer periods)

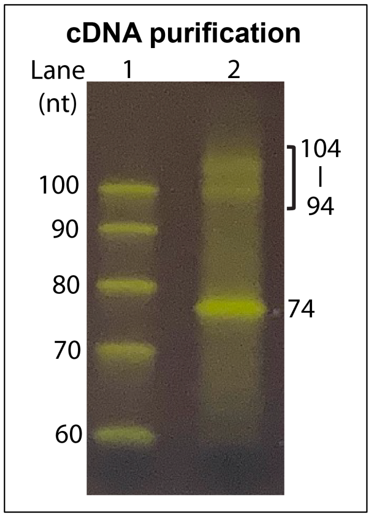


**G. Circularization of cDNA**

81. Prepare circularization master mix on ice:

| | Volume (µL) | Final |
|---|---|---|
| 10x CircLigase buffer | 2 | 1x |
| 1 mM ATP | 1 | 50 µM |
| 50 mM MnCl2 | 1 | 2.5 mM |
| CircLigase (100 U/µL) | 1 | 100U |

82. Add 5 µL of circularization master mix to each sample and mix. Now the sample volume is 20 µL.
83. Incubate with the following program in a thermocycler:
    60°C – 2 hr
    80°C – 10 min
    4°C – hold.
    (Optional stopping point: store at -20°C)

### H. qPCR quantification of circularized cDNA

84. Prepare a dilution series of the positive control:
    - Mix 2 µL of a **1 µM** stock with 198 µL NF-H2O (10 nM).
    - Mix 10.2 µL of the 10 nM solution above with 89.8 µL of NF-H2O (1.02 nM).
    - Serially dilute 3 µL of the stock into 9 µL of NF-H2O to prepare a 1:4 dilution (~256 pM), a 1:16 dilution (64 pM), a 1:64 dilution (16 pM), a 1:256 dilution (4 pM), and a 1:1024 dilution (1 pM).
    - **Also prepare 0.5 nM negative control (reverse transcription primer).**
85. Mix 1 µL of circularized cDNA from Step 83 with 9 µL NF-H2O.
86. Set up qPCR as follows:
    - Serial dilutions and controls (total of 8): 1.02 nM, 256 pM, 64 pM, 16 pM, 4 pM, 1 pM, NF-H2O, negative control.
    - Each circularization requires 2 technical replicates.
87. Prepare qPCR master mix on ice:

| | 1 reaction | ________ reactions | Final |
|---|---|---|---|
| NF-H2O | 6.4 | | |
| 2x Luna qPCR master mix | 10 | | 1X |
| 10 µM primer F | 0.8 | | 0.4 µM |
| 10 µM primer R | 0.8 | | 0.4 µM |

88. For each reaction, combine 18 µL of qPCR master mix with 2 µL of template, pipette 10 times.
89. Spin down and perform qPCR amplification using the following cycling conditions:
    - 95° C, 60 s
    - Repeat 40 cycles of
      95 °C, 15 s
      59 °C, 30 s
    - Melting curve analysis.

90. Fit a standard curve to the Cq values for the serial dilution series. The 1.02 nM sample should have a Cq of roughly 8. Verify that the negative control and the blank reactions have Cq values much higher than the standard curve or circularized cDNA samples.
91. Determine the template concentration in the circularization reactions based on the standard curve.

92. Select the amount of template and the number of cycles for the library construction PCR based on the table below for a 50-μL PCR reaction.

| Template concentration | Cycles needed |
|---|---|
| 800 pM | 7 |
| 400 pM | 8 |
| 200 pM | 9 |
| 100 pM | 10 |
| 50 pM | 11 |
| 25 pM | 12 |
| 12.5 pM | 13 |
| 6.25 pM | 14 |
| 3.125 pM | 15 |
| 1.6 pM | 16 |

93. If a template is at 700 pM, 2.3 μL of template will have a concentration of 32 pM in a 50 μL reaction, so ~12 cycles will be needed.
    - Make sure the template comprises no more than 10% of the final PCR volume and ideally no more than 5%. This helps minimize reannealed duplexes.

**I. Library construction PCR**

94. For each sample, set up a 50-μL PCR reaction on ice:

| | Volume (μL) |
|---|---|
| NF-H2O | 20-X |
| 2X Phusion HF master mix | 25 |
| 10 μM Forward library PCR primer | 2.5 |
| 10 μM Reverse ***INDEXED*** primer | 2.5 |
| Circularized cDNA | X |

95. Purify the PCR products using a DNA Clean & Concentrator 5 kit. Elute in 25.5 μL NF-H2O (recover ~25 μL).
96. Add 5 μL of 6x DNA gel loading dye (purple) to each sample.
97. Prepare the 20-bp ladder.

98. Set up a pre-cast 8% polyacrylamide non-denaturing gel (prepare 450 mL 1x TBE and save 40 mL on ice for staining).

99. Load 3 adjacent wells with 10 µL each of the purified PCR samples; separate different samples with 3 µL of 20 bp ladder.

100. Run the gel at 200 V for 40 min in 1X TBE.

101. Stain the gel with ice-cold 1x TBE/SYBR Gold for 3 min (4 µL SYBR-GOLD in 40 mL TBE buffer above)

102. Excise the expected library band:
   - For 20-30-nt footprints, the library should be **162-172 bp**.
   - Avoid bands that are ~145 bp and below (these are products resulting from self-ligated universal miRNA cloning linker and self-circularized reverse transcription primer).

103. Extract DNA as described in Step 79.

104. Resuspend the DNA in 11 µL of 10 mM Tris pH 8.

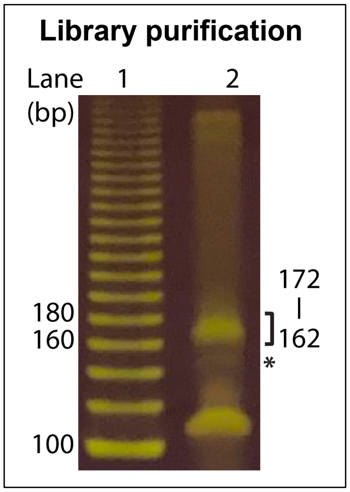


### J. Library QC and pooling

105. Quantify the library DNA with a Qubit DNA HS assay.

106. Analyze library size distributions with Agilent Fragment Analyzer. Libraries should be roughly 170 bp.

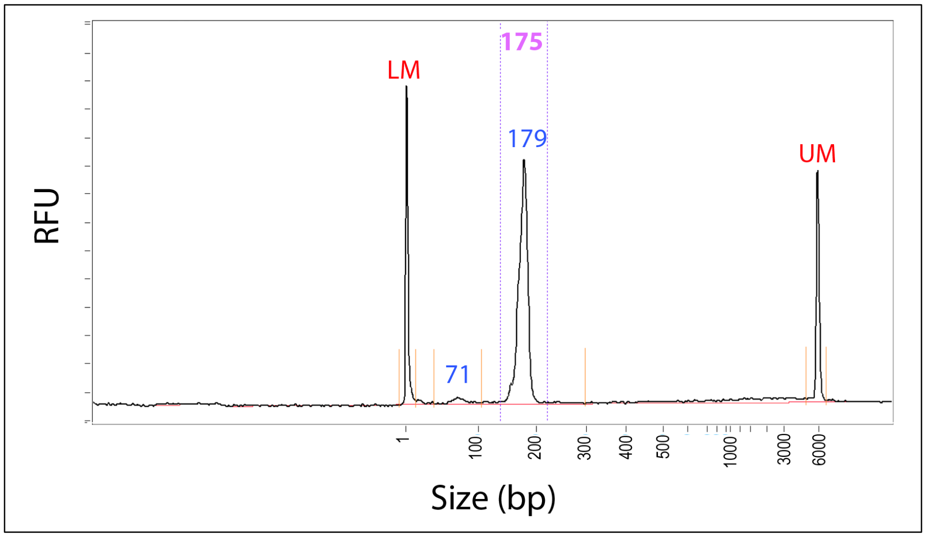


107. Pool libraries with the same molarity.

108. Proceed with single-end 50-bp sequencing.
