## Supplemental File S4 for "A custom library construction method for super-resolution ribosome profiling in Arabidopsis"

#Step 1 code (FASTQC for quality check):
#Run in linux server or cluster
#$OUTPUT output directory
#The code below is to check all .gz file in the directory

fastqc -o $OUTPUT -t 10 *.gz

#Step 2 code (FASTX toolkit to trim adaptor sequence):
#Run in linux server or cluster
#$INPUT input directory
#$OUTPUT output directory

zcat $INPUT/NEB123.fastq.gz | fastx_clipper -o $OUTPUT/NEB123.trim.fastq -a CTGTAGGCACCATCAAT -l 20 -c -n -v -Q33


#Step 3 code:
#Run in linux server or cluster
#$CF is the directory contain FASTA files
#$Contam5 is the directory with the index for the contamination sequences 
#$OUTPUT output directory

bowtie2-build $CF/Araport11_201606_rRNA.fasta,$CF/Araport11_201606_tRNA.fasta,$CF/Araport11_201606_snRNA.fasta,$CF/Araport11_201606_snoRNA.fasta,$CF2/AT3G06365_AT2G03875.fa Contam5

bowtie2 -L 20 -p 8 -x $Contam5 $OUTPUT/NEB123.trim.fastq --un-gz $OUTPUT/NEB123.noContam5.fastq.gz

#Step 4 A code:
#Run in linux server or cluster
#$starIndex1 is the directory contain FASTA files
#FASTA is the path to the TAIR10 genome FASTA file
#GTF is the path to the Araport11 GTF file 
#$INPUT is the directory with the index for the contamination sequences 
#$OUTPUT is the output directory


STAR --runThreadN 12 \
--runMode genomeGenerate \
--genomeDir $starIndex1 \
--genomeFastaFiles $FASTA \
--sjdbGTFfile $GTF \
--sjdbOverhang 34 \

#Step 4 B code:
#Run in linux server or cluster
#$starIndex1 is the directory contain FASTA files
#$INPUT is the directory with the NEB123.noContam5.fastq.gz file
#$OUTPUT is the output directory


STAR --runThreadN 10 \
--genomeDir $starIndex1 \
--readFilesCommand zcat \
--readFilesIn $INPUT/NEB123.noContam5.fastq.gz \
--alignIntronMax 5000 \
--alignIntronMin 15 \
--outFilterMismatchNmax 1 \
--outFilterMultimapNmax 20 \
--outFilterType BySJout \
--alignSJoverhangMin 8 \
--alignSJDBoverhangMin 2 \
--outSAMtype BAM SortedByCoordinate \
--quantMode TranscriptomeSAM \
--outSAMmultNmax 1 \
--outMultimapperOrder Random \
--outFileNamePrefix "star_ribo_NEB123" \

#Step 5 code:
#Create 2bit files
#Run the code in R (v. 4.1.3) locally, on server or cluster

library(Biostrings)
library(rtracklayer)
library(RiboseQC)
At_genome_seqs <- Biostrings::readDNAStringSet("~/Desktop/New_Riboseq/TAIR10_chr_all_2.fas") 
At_genome_seqs <- replaceAmbiguities(At_genome_seqs)
test_2bit_out <- file.path("~/Desktop/New_Riboseq/", "At.2bit") #Do not contain Chr
#~/Desktop/ORFquant/At_chr.2bit contain Chr
rtracklayer::export.2bit(At_genome_seqs, test_2bit_out)

setwd("~/Desktop/New_Riboseq")
prepare_annotation_files(annotation_directory = ".",
                         twobit_file = "~/Desktop/New_Riboseq/At.2bit",
                         gtf_file = "~/Desktop/New_Riboseq/Araport11_20220629.gtf",scientific_name = "Arabidopsis.thaliana",
                         annotation_name = "TAIR10",export_bed_tables_TxDb = F,forge_BSgenome = T,create_TxDb = T)

RiboseQC_analysis(annotation_file="~/Desktop/New_Riboseq/Araport11_20220629.gtf_Rannot",bam_files = "~/Desktop/New_Riboseq/Contam4/star_ribo_NEB123_Contam4_Aligned.sortedByCoord.out.bam",report_file = "NEB123_noContam4.html",write_tmp_files = F)

Example gtf file (the 9th column is the key for running RiboseQC):
1	Araport11	gene	3631	5899	.	+	.	gene_id "AT1G01010"; gene_biotype "protein_coding";
1	Araport11	CDS	3760	3913	.	+	0	gene_id "AT1G01010"; transcript_id "AT1G01010.1"; gene_biotype "protein_coding";
1	Araport11	CDS	3996	4276	.	+	2	gene_id "AT1G01010"; transcript_id "AT1G01010.1"; gene_biotype "protein_coding";
1	Araport11	CDS	4486	4605	.	+	0	gene_id "AT1G01010"; transcript_id "AT1G01010.1"; gene_biotype "protein_coding";
1	Araport11	CDS	4706	5095	.	+	0	gene_id "AT1G01010"; transcript_id "AT1G01010.1"; gene_biotype "protein_coding";
1	Araport11	CDS	5174	5326	.	+	0	gene_id "AT1G01010"; transcript_id "AT1G01010.1"; gene_biotype "protein_coding";
1	Araport11	CDS	5439	5630	.	+	0	gene_id "AT1G01010"; transcript_id "AT1G01010.1"; gene_biotype "protein_coding";
1	Araport11	transcript	3631	5899	.	+	.	gene_id "AT1G01010"; transcript_id "AT1G01010.1"; gene_biotype "protein_coding";
1	Araport11	exon	3631	3913	.	+	.	gene_id "AT1G01010"; transcript_id "AT1G01010.1"; gene_biotype "protein_coding";
1	Araport11	exon	3996	4276	.	+	.	gene_id "AT1G01010"; transcript_id "AT1G01010.1"; gene_biotype "protein_coding";
1	Araport11	exon	4486	4605	.	+	.	gene_id "AT1G01010"; transcript_id "AT1G01010.1"; gene_biotype "protein_coding";
1	Araport11	exon	4706	5095	.	+	.	gene_id "AT1G01010"; transcript_id "AT1G01010.1"; gene_biotype "protein_coding";
1	Araport11	exon	5174	5326	.	+	.	gene_id "AT1G01010"; transcript_id "AT1G01010.1"; gene_biotype "protein_coding";
1	Araport11	exon	5439	5899	.	+	.	gene_id "AT1G01010"; transcript_id "AT1G01010.1"; gene_biotype "protein_coding";
1	Araport11	five_prime_UTR	3631	3759	.	+	.	gene_id "AT1G01010"; transcript_id "AT1G01010.1"; gene_biotype "protein_coding";
1	Araport11	three_prime_UTR	5631	5899	.	+	.	gene_id "AT1G01010"; transcript_id "AT1G01010.1"; gene_biotype "protein_coding";
1	Araport11	gene	163278	166353	.	+	.	gene_id "AT1G01448"; gene_biotype "antisense_long_noncoding_rna";
1	Araport11	transcript	163278	166353	.	+	.	gene_id "AT1G01448"; transcript_id "AT1G01448.1"; gene_biotype "antisense_long_noncoding_rna";
1	Araport11	exon	163278	163516	.	+	.	gene_id "AT1G01448"; transcript_id "AT1G01448.1"; gene_biotype "antisense_long_noncoding_rna";
1	Araport11	exon	163934	164103	.	+	.	gene_id "AT1G01448"; transcript_id "AT1G01448.1"; gene_biotype "antisense_long_noncoding_rna";
1	Araport11	exon	164225	164686	.	+	.	gene_id "AT1G01448"; transcript_id "AT1G01448.1"; gene_biotype "antisense_long_noncoding_rna";
1	Araport11	exon	164771	165380	.	+	.	gene_id "AT1G01448"; transcript_id "AT1G01448.1"; gene_biotype "antisense_long_noncoding_rna";
1	Araport11	exon	165449	166353	.	+	.	gene_id "AT1G01448"; transcript_id "AT1G01448.1"; gene_biotype "antisense_long_noncoding_rna";
1	Araport11	transcript	163278	166353	.	+	.	gene_id "AT1G01448"; transcript_id "AT1G01448.2"; gene_biotype "antisense_long_noncoding_rna";
1	Araport11	exon	163278	163516	.	+	.	gene_id "AT1G01448"; transcript_id "AT1G01448.2"; gene_biotype "antisense_long_noncoding_rna";
1	Araport11	exon	163934	164103	.	+	.	gene_id "AT1G01448"; transcript_id "AT1G01448.2"; gene_biotype "antisense_long_noncoding_rna";
1	Araport11	exon	164225	164686	.	+	.	gene_id "AT1G01448"; transcript_id "AT1G01448.2"; gene_biotype "antisense_long_noncoding_rna";
1	Araport11	exon	164771	165380	.	+	.	gene_id "AT1G01448"; transcript_id "AT1G01448.2"; gene_biotype "antisense_long_noncoding_rna";
1	Araport11	exon	165449	165656	.	+	.	gene_id "AT1G01448"; transcript_id "AT1G01448.2"; gene_biotype "antisense_long_noncoding_rna";
1	Araport11	exon	165769	166353	.	+	.	gene_id "AT1G01448"; transcript_id "AT1G01448.2"; gene_biotype "antisense_long_noncoding_rna";
1	Araport11	transcript	163302	166307	.	+	.	gene_id "AT1G01448"; transcript_id "AT1G01448.3"; gene_biotype "antisense_long_noncoding_rna";
1	Araport11	exon	163302	163516	.	+	.	gene_id "AT1G01448"; transcript_id "AT1G01448.3"; gene_biotype "antisense_long_noncoding_rna";
1	Araport11	exon	163934	164103	.	+	.	gene_id "AT1G01448"; transcript_id "AT1G01448.3"; gene_biotype "antisense_long_noncoding_rna";
1	Araport11	exon	164225	164686	.	+	.	gene_id "AT1G01448"; transcript_id "AT1G01448.3"; gene_biotype "antisense_long_noncoding_rna";
1	Araport11	exon	164771	165380	.	+	.	gene_id "AT1G01448"; transcript_id "AT1G01448.3"; gene_biotype "antisense_long_noncoding_rna";
1	Araport11	exon	165449	165656	.	+	.	gene_id "AT1G01448"; transcript_id "AT1G01448.3"; gene_biotype "antisense_long_noncoding_rna";
1	Araport11	exon	165983	166307	.	+	.	gene_id "AT1G01448"; transcript_id "AT1G01448.3"; gene_biotype "antisense_long_noncoding_rna";
Pt	Araport11	gene	60741	61430	.	+	.	gene_id "ATCG00530"; gene_biotype "protein_coding";
Pt	Araport11	CDS	60741	61430	.	+	0	gene_id "ATCG00530"; transcript_id "ATCG00530.1"; gene_biotype "protein_coding";
Pt	Araport11	transcript	60741	61430	.	+	.	gene_id "ATCG00530"; transcript_id "ATCG00530.1"; gene_biotype "protein_coding";
Pt	Araport11	exon	60741	61430	.	+	.	gene_id "ATCG00530"; transcript_id "ATCG00530.1"; gene_biotype "protein_coding";
Mt	Araport11	gene	366086	366700	.	-	.	gene_id "ATMG01410"; gene_biotype "protein_coding";
Mt	Araport11	CDS	366086	366700	.	-	0	gene_id "ATMG01410"; transcript_id "ATMG01410.1"; gene_biotype "protein_coding";
Mt	Araport11	transcript	366086	366700	.	-	.	gene_id "ATMG01410"; transcript_id "ATMG01410.1"; gene_biotype "protein_coding";
Mt	Araport11	exon	366086	366700	.	-	.	gene_id "ATMG01410"; transcript_id "ATMG01410.1"; gene_biotype "protein_coding";

Step 6A: Kallisto (v0.46.1) indexing:

#$OUTPUT is the path to the output folder
#$FASTA is the path to the transcript fasta file
kallisto index -i $OUTPUT/transcripts.idx $FASTA -k 19

Step 6B: Kallisto mapping for Ribo-seq reads
# The three fastq.gz files are from the three tech replicates using step 1 and step 2 code to analyze 

kallisto quant -i $Index/transcripts.idx -o $OUTPUT1 -t 10 --single -l 28 -s 2 $INPUT/NEB1.noContam5.fastq.gz
kallisto quant -i $Index/transcripts.idx -o $OUTPUT2 -t 10 --single -l 28 -s 2 $INPUT/NEB2.noContam5.fastq.gz
kallisto quant -i $Index/transcripts.idx -o $OUTPUT3 -t 10 --single -l 28 -s 2 $INPUT/NEB3.noContam5.fastq.gz

Step 6C: Correlation plot
library(dplyr)
library(corrplot)
RiboD1 <- read.delim("~/Desktop/New_Riboseq/NEB1_abundance.tsv",header=T,sep="\t",stringsAsFactors = F,quote = "")
RiboD2 <- read.delim("~/Desktop/New_Riboseq/NEB2_abundance.tsv",header=T,sep="\t",stringsAsFactors = F,quote = "")
RiboD3 <- read.delim("~/Desktop/New_Riboseq/NEB3_abundance.tsv",header=T,sep="\t",stringsAsFactors = F,quote = "")

TPM=data.frame(RiboD1=RiboD1$tpm,RiboD2=RiboD2$tpm,RiboD3=RiboD3$tpm)

TPM$Ribo_mean <-rowMeans(TPM[,1:3])

TPM2 <- TPM %>% filter(Ribo_mean>0.1) %>% filter(Ribo_mean<200)

R2 <- round(cor(TPM2[,1:3]),2)

pdf("~/Desktop/New_Riboseq/NEB_samples_correlation.pdf",width =4,height = 4)
corrplot(R2, method = "number",type="upper")
dev.off()

Step 7: Calculate 3-nt periodicity
# Load the results_RiboseQC file from Ribo-seQC output
load("~/Desktop/New_Riboseq/star_ribo_NEB123Aligned.sortedByCoord.out.bam_results_RiboseQC")
# Extract P_sites_subcodon
P_sites_subcodon_readCount = res_all[["profiles_P_sites"]][["P_sites_subcodon"]][["nucl"]][["28"]]
# Print out P_sites_subcodon_readCount
P_sites_subcodon_readCount

# Get the CDS region
P_CDS = P_sites_subcodon_readCount[51:149] #93 nt total, 33 nt start and after, 33 nt in the middle, 27 nt from -2 codon of stop codon to -11 codon of stop codon

# Get total number of reads for 3 frames
F1 = sum(P_CDS[seq(1,93,by=3)])
F2 = sum(P_CDS[seq(2,93,by=3)])
F3 = sum(P_CDS[seq(3,93,by=3)])
F1/(F1+F2+F3)*100 #[1] 91.03305
